# Comparison of materials for rapid passive collection of environmental DNA

**DOI:** 10.1101/2021.11.08.467842

**Authors:** Cindy Bessey, Yuan Gao, Yen Bach Truong, Haylea Miller, Simon Neil Jarman, Oliver Berry

## Abstract

Passive collection is an emerging sampling method for environmental DNA (eDNA) in aquatic systems. Passive eDNA collection is inexpensive, efficient, and requires minimal equipment, making it suited to high density sampling and remote deployment. Here, we compare the effectiveness of nine membrane materials for passively collecting fish eDNA from a 3 million litre marine mesocosm. We submerged materials (cellulose, cellulose with 1% and 3% chitosan, cellulose overlayed with electrospun nanofibers and 1% chitosan, cotton fibres, hemp fibres and sponge with either zeolite or active carbon) for intervals between five and 1080 minutes. We show that for most materials, with as little as five minutes submersion, mitochondrial fish eDNA measured with qPCR, and fish species richness measured with metabarcoding, was comparable to that collected by conventional filtering. Furthermore, PCR template DNA concentrations and species richness were generally not improved significantly by longer submersion. Species richness detected for all materials ranged between 11 to 37 species, with a median of 27, which was comparable to the range for filtered eDNA (19-32). Using scanning electron microscopy, we visualised biological matter adhered to the surface of materials, rather than entrapped, with images also revealing a diversity in size and structure of putative eDNA particles.

Environmental DNA can be collected rapidly from seawater with a passive approach and using a variety of materials. This will suit cost and time-sensitive biological surveys, and where access to equipment is limited.

## Introduction

Environmental DNA metabarcoding is a sensitive, non-invasive and broadly applicable tool for species detection, including biodiversity measurement and biosecurity surveillance (Taberlet et al. 2018, Deiner et al. 2017). Macro-organisms shed their DNA into the air, soil, and water, which can be sampled by collecting, extracting, amplifying, sequencing, and ultimately identified by comparing against a reference database of known DNA sequences (Taberlet et al. 2018, Thomsen et al. 2015). Diverse applications have been developed for eDNA metabarcoding and the field has grown rapidly in recent years (Koziol et al. 2019, Jarman et al. 2018). Nevertheless, a common limitation of eDNA studies is a lack of replication (Buxton et al. 2021, Derocles et al. 2018, Dickie et al. 2018, Zinger et al. 2017). For aquatic systems this is in part due to the logistical challenge of filtering sufficient water (Bessey et al. 2020).

Passive collection methods (Bessey et al, 2021, Kirtane et al. 2020), which involves direct submersion into a water body of materials that collect eDNA, facilitate increased replication because they are cheaper, simpler, and faster to apply than active filtration. This enables analyses that are generally not practical with water filtering as an eDNA collection method. Frequency of occurrence methods become more feasible, as well as mapping residence of species of interest. For studies investigating diversity, greater biological replication improves the reliability of both alpha and beta diversity estimates (Zinger et al. 2017, Prosser 2010). Furthermore, because passive eDNA requires minimal or no supporting technology it suits deployment to remote environments, and by non-experts.

Few studies investigate the mechanisms or optimal material properties needed for passive eDNA collection. Kirtane et al. (2020) used adsorbent-filled sachets of Montmorillonite clay and granular activated carbon to passively capture eDNA in freshwater laboratory, microcosm and field experiments. In the laboratory, they found that extracellular DNA adsorbed to these materials at different rates, depending on the water matrix. In their field experiments, granular activated carbon sachets captured significantly more eDNA than clay and detected as many fish species as a 1 L conventional grab sample. These materials were chosen for their high adsorption capacity to trap DNA but also for their low adsorption affinity to allow high yield during extraction. They suggest adsorption mechanisms for granular activated carbon are dependent on the water matrix, whereas that of clay is more dependent on adsorption kinetics and capacity. Bessey et al. (2021) compared the effectiveness of positively charged nylon and non-charged cellulose ester membrane materials for passive collection of fish eDNA at both a species-rich tropical and species-poor temperate marine site. They found that both materials detected fish as effectively as conventional active eDNA filtration methods in temperate systems and provided similar estimates of total fish biodiversity but differed in tropical waters. Their materials were chosen to investigate the possible role of electrostatic attraction and because both are commonly used in conventional aquatic eDNA studies using filtration methods. The observations that significant material effects exist, and may be system specific, indicates there is potential for improvements to passive eDNA collection through material selection that could create greater efficiencies for users.

The optimal submersion time for efficient passive eDNA collection is also unclear. Kirtane et al. (2020) found that, regardless of material used (clay or granular carbon) or water matrix (molecular grade water, microcosm tank water, or natural creek water), an equilibrium concentration of eDNA was absorbed in less than 24 hours. In field trials, they also found that fish species detection did not significantly increase with longer submersion duration (7 days compared to 21). In both tropical and marine waters, Bessey et al. (2021) likewise found that increased submersion time did not increase species richness (comparing 4, 8, 12, and 24 hours of submersion). Combined, these studies indicate that long-duration submersion (days or hours) may not be necessary and therefore, investigations into minimal submersion times are another potential avenue to increase passive eDNA collection efficiency.

Using a DNA metabarcoding approach, here we evaluate the effect of materials and submersion time on the efficiency with which fish eDNA could be collected passively from a large marine mesocosm. We also use scanning electron microscopy to visualise modes of eDNA adherence or entanglement to the different materials. We show that for most materials, passively collected eDNA consistently performs similarly to conventionally filtered eDNA samples, and that high collection efficiency can be achieved in as little as five minutes.

## Materials and Methods

### Study Site and Design

Sampling was conducted in the main tank at The Aquarium of Western Australia (AQWA; aqwa.com.au), which offered a relatively controlled system containing 50 known fish species in three million litres of seawater. This system draws incoming seawater from 0.5 m below the seabed (natural sand filter) of the nearshore ocean waters. It is then filtered (pressure glass media filter) before entering the AQWA facility where the water supplies several display tanks before entering the main tank of the mesocosm. The main tank has its own gravity filter system (volume of filter tank is 2 million litres) that uses a 50 cm sand bed with 2 mm (± 0.5 mm) size particles, over 50 cm of 6 mm (± 3 mm) gravel. The turnover rate between the gravity filter and main tank is 5 million litres every 2 hours. Passive eDNA sampling was conducted between 8am and 4pm on January 21 and 22, 2021, by submerging nine different membrane materials just below the surface in the mesh pockets of a pearl oyster aquaculture frame (Fig.1, see Bessey et al. 2021). Each of the nine membrane materials were deployed in quadruplicate for specified time intervals (5, 10, 30, 60 minutes and overnight for 18 hours) to examine whether increased submersion time led to increased eDNA collection. Of the quadruplicate samples, three were used for eDNA extractions while the other was used for scanning electron microscopy to visualise how eDNA collected on the different membrane surfaces.

**Figure 1.**
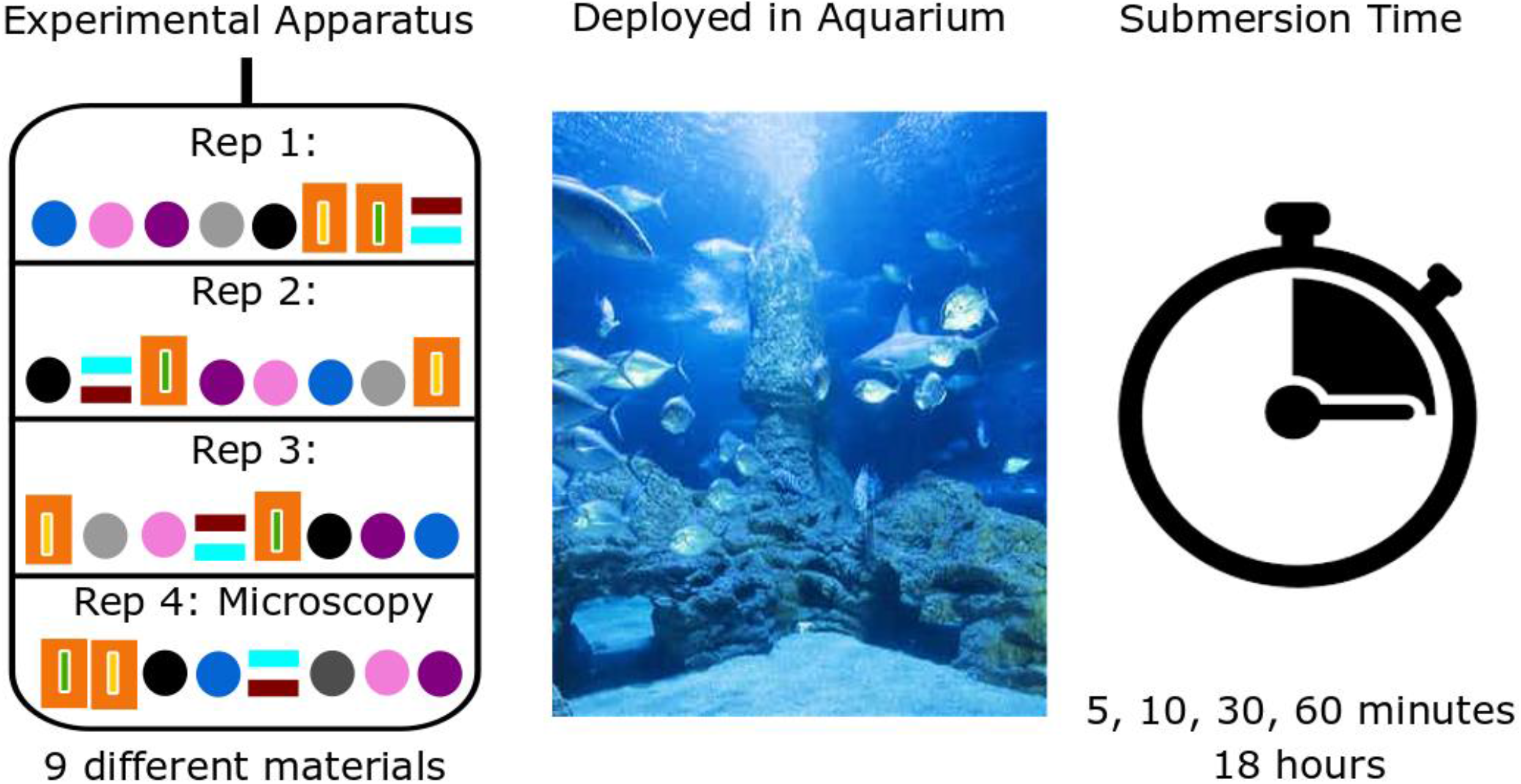
Experimental design for testing passive eDNA collection with nine different membrane materials in a controlled mesocosm setting with varying soak times.

### Membrane Materials

We trialled nine different membrane materials (Table 1). The first was a cellulose ester membrane (0.45 μm Pall GN-6 Metricel®) commonly used in eDNA studies (Tsuji et al. 2019). To investigate whether chitosan coating would increase eDNA capture, the cellulose membranes were impregnated with either 1% or 3% chitosan (w/w) which was then crosslinked under glutaraldehyde vapour to confer stability. Loadings of chitosan on the membranes were confirmed by Fourier-transform infrared spectroscopy (FT-IR) as well as by staining with the anionic dye Eosin Y. Chitosan is a polycation polymer that efficiently binds anionic DNA under acidic conditions and has been used for DNA enrichment and purification (Pandit et al. 2015). Chitosan is derived from chitin in crustacean shells and is readily available, inexpensive and biocompatible. To investigate if eDNA would become entrapped in highly complex materials, we trialled overlaying the cellulose esters with electrospun nanofibres, while also trialling a combination of electrospun nanofibers that were subsequently covered in a 1% chitosan (w/w). Electrospinning is a technique for producing fibres from submicron down to nanometer in diameter with high surface area (Bhardwaj and Kundu 2010). We used solution electrospinning, where the polymer(s) and other additive materials are firstly dissolved in a suitable solvent at an optimized concentration before electrospinning. A high Voltage electric field is applied to the droplet of fluid coming out of the tip of a die or spinneret, which acts as one of the electrodes. When the electric field supply is strong enough, it will lead to the droplet formation and finally to the ejection of a charged jet from the tip of the cone accelerating toward the counter collector electrode leading to the formation of a nanofibrous membrane. These nanofibrous membranes have found applications in many areas, including biomedical areas (e.g., scaffolds for tissue engineering, drug delivery, wound dressing, and medical implants), filtration, protective textiles, and battery cells (Gao et al. 2014). Our electrospinning was carried out using polyether based thermoplastic polyurethane (TPU) grade (RE-FLEX® 585A, Townsend Chemicals) with a 10% w/v solution in dimethyl formamide solvent (DMF) using a 23 G needle spinneret, with an applied voltage of 20kV at 15 cm from the collecting drum. To ensure sufficient physical robustness for use in the marine environment, a composite was prepared using a thermal bonding [Protechnic 114P (13 gsm)] net material to bond the electrospun membrane attached through thermal adhesive. This backing plate was needed to prevent the nanofibre cellulose membranes from curling, and therefore, we also trialled these backing plates separately in the downstream processing to determine their effect on eDNA capture. We also trialled natural fibres, cotton and hemp, which were contained in a nylon bag for practical deployment purposes so they would remain anchored within the mesh of the pearl frame. A subset of nylon bags was retained for downstream processing in the same fashion as the trialled materials. These cotton and hemp fibres were 5 mm in diameter and cut into 40 mm lengths so they could fit in a 2 mL Eppendorf tube for DNA extraction. Finally, we trialled two sponge materials that would be highly robust in aquatic settings: one was a tightly woven filter pad with 100% active carbon (Aqua One®), while the other was a tightly woven filter pad with zeolite (Aqua One®). The sponge was cut into 40 mm rectangular lengths and had a 5 mm width and depth. All materials were placed under ultra-violet sterilizing light for a minimum of 30 min, except for the cellulose membranes which were certified sterile upon purchase.

**Table 1.**
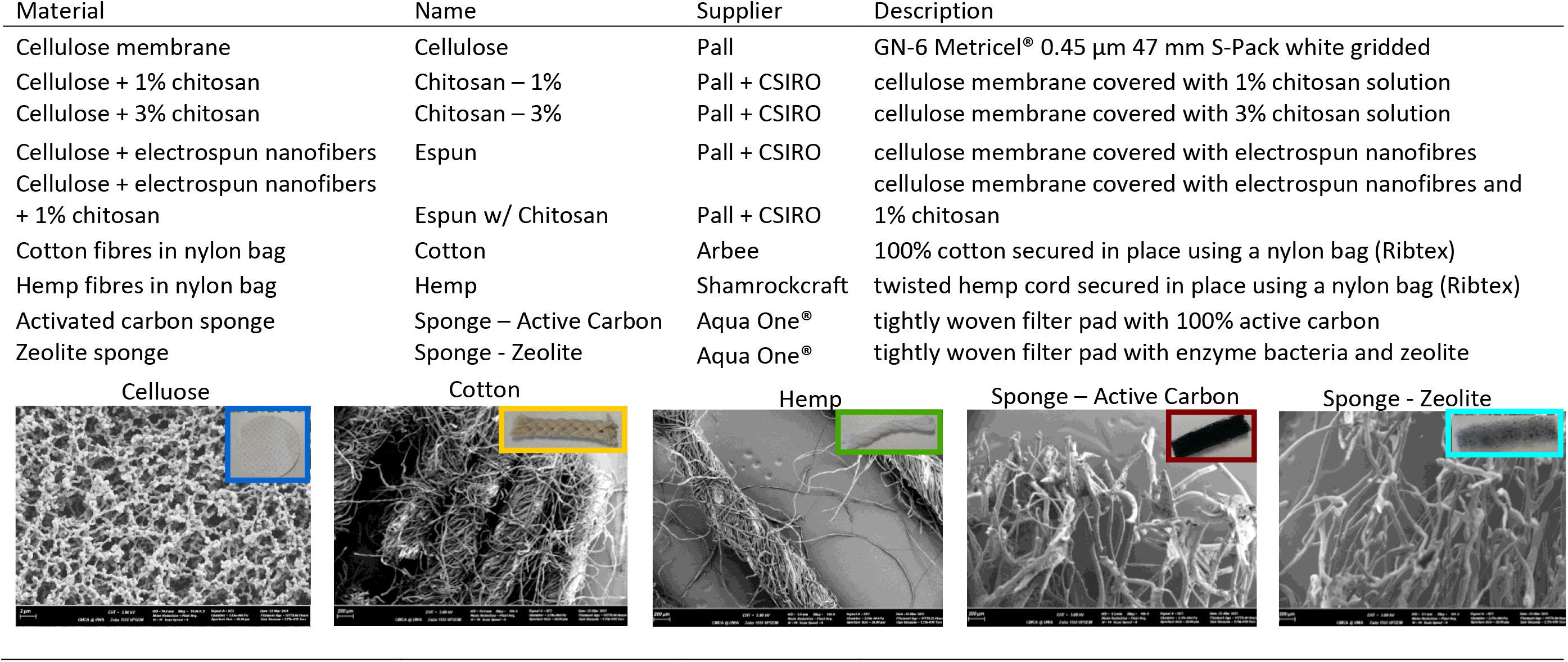
Materials trialled for passive eDNA collection accompanied by scanning electron microscopy pictures of each material at 10,000 × (Cellulose) and 100 × (all other materials) magnification, inlaid with pictures of the materials prior to deployment.

### Scanning Electron Microscopy

We used scanning electron microscopy (SEM) to qualitatively investigate how biological matter attaches to each of the different materials. Scanning electron microscopy uses a focused beam of high-energy electrons to generate a variety of signals at the surface of solid specimens. These signals are converted into 2-dimensional high-resolution images and reveal information about the external morphology (texture), chemical composition, and crystalline structure and orientation of materials making up the sample. A subsection of each membrane material was dissected using sterile surgical scissors and mounted onto 10 mm dimeter SEM stubs with an adhesive carbon tab to prevent charge build-up. The stubs were then air dried at ambient temperature in a fume hood while partially covered to prevent dust or debris on the samples. Once completely dry, samples were coated with a thin layer of platinum (sputter coated with 3 nm of platinum using a Leica EM MED020; Leica Microsystems, Inc. Buffalo Grove, IL); a fine conducting material for high resolution electron imaging. The samples were visualized and imaged on a Zeiss 1555 VP-FESEM with SmartSEM software (Zeiss, Germany) at the Centre for Microscopy, Characterisation and Analysis (CMCA), University of Western Australia, Perth, Western Australia. We provide example SEM images of materials at 10,000 × (cellulose) and 100 × (cotton, hemp, sponge – active carbon and sponge - nitrate) magnification prior to deployment (Table 1) and provide an example of all deployed materials with biological matter attached. Due to limited supplies of chitosan and electrospun nanofiber covered cellulose membranes, none were available for SEM imagining prior to deployment.

### Active eDNA Collection

We collected water for active eDNA filtration to compare to the results of passive eDNA collection. Five 1 L surface water samples were collected in sterile 1 L containers at five different times over the day and filtered with cellulose ester membranes (47 mm diameter, 0.45 μm pore size) using a peristaltic Sentino® Microbiology Pump on a clean benchtop at the aquarium facility. All water samples were taken on the first day.

### Contamination control

Sterile technique was used throughout the experiment and consisted of wearing gloves and using sterile tweezers to handle all materials. All materials were frozen after collection and stored at −20°C until further processing in the laboratory. All collection and deployment apparatus were sterilized by soaking in 10% bleach solution for at least 15 minutes and rinsed in deionized water.

### eDNA extraction from passive collection materials

All cellulose ester materials, as well as the nylon bags, were cut or flash frozen (−80°C) and crushed into small pieces that were placed in a 2 mL Eppendorf tube in preparation for extraction. All other materials were placed directly into a 2mL Eppendorf tube as is for extraction. Total nucleic acid was extracted from all materials in the same fashion using a DNeasy Blood and Tissue Kit (Qiagen; Venlo, Netherlands), with an additional 40 μL of Proteinase K used during a three-hour digestion period at 56°C on rotation (300 rpm). DNA was eluted into 200 μL AE buffer. All extractions took place in a dedicated DNA extraction laboratory using a QIAcube (Qiagen; Venlo, Netherlands), where benches and equipment were routinely bleached and cleaned.

### DNA metabarcode amplification for fish detection

We followed the same procedures used by Bessey et al. (2021). One-step quantitative polymerase chain reactions (qPCR) were performed in duplicate for each sample using 2 μL of extracted DNA and a mitochondrial DNA 16S rDNA universal primer set targeting fish taxa (16SF/D 5ʹ GACCCTATGGAGCTTTAGAC 3ʹ and 16S2R-degenerate 5ʹ CGCTGTTATCCCTADRGTAACT 3ʹ; Berry et al. 2017, Deagle et al. 2007), with the addition of fusion tag primers unique to each sample that included Illumina P5 and P7 adaptors. A single round of qPCR was performed in a dedicated PCR laboratory. Quantitative PCR reagents were combined in a dedicated clean room and included 5 μL AllTaq PCR Buffer (QIAGEN; Venlo, Netherlands), 0.5 μL AllTaq DNA Polymerase, 0.5 μL dNTPs (10 mM), 1.0 μL Ultra BSA (500 μg/μL), SYBR Green I (10 units/μL), 0.5 μL forward primer (20 μM) and 5.0 μL reverse primer (20 μM), 2 μL of DNA and Ultrapure™Distilled Water (Life Technologies) made up to 25 μL total volume. Mastermix was dispensed manually and qPCR was performed on a CFX96 Touch™ Real-Time PCR Detection System (Bio-Rad, California, USA) using the following conditions: initial denaturation at 95°C for 5 min, followed by 40 cycles of 30 s at 95°C, 30 s at the primer annealing temperature 54°C, and 45 s at 72°C, with a final extension for 10 min at 72°C. All duplicate qPCR products from the same subsample were combined prior to library pooling. The mean Cq value from qPCR duplicates was used as an indication of initial DNA copy number. A sequencing library was made by pooling amplicons into equimolar ratios based on qPCR Ct values and sequenced on an Illumina Miseq platform (Illumina; San Diego, USA). The libraries were size selected using a Pippin Prep (Sage Science, Beverly, USA) and purified using the Qiaquick PCR Purification Kit (Qiagen; Venlo, Netherlands). The volume of purified library added to the sequencing run was determined by quantifying the concentration (Murray et al. 2015) using a Qubit 4 fluorometer (ThermoFisher Scientific). The library was unidirectionally sequenced using a 300 cycle MiSeq® V2 Reagent Kit and standard flow cell.

PCR plates included blank laboratory extraction controls (extraction reagents used with no DNA template), PCR negative controls (2 μL of DI water used rather than DNA template) and positive controls (dhufish; *Glaucosoma hebraicum* and swordfish; *Xiphias gladius*). Dhufish inhabit the mesocosm, whereas swordfish do not, so the latter was a more appropriate positive control. No negative control (extraction or PCR) contained more than 17 reads, with the maximum number of reads per fish species being four. Therefore, we used a detection rate of greater than five sequences to classify something as a positive detection. All positive controls amplified multiple reads identifying dhufish and swordfish with 100% identity. Swordfish was not detected in any sample except for our positive PCR control.

### DNA sequence data processing

Our DNA sequence data processing is detailed in Bessey et al. (2021), it directly follows the procedure described at https://pythonhosted.org/OBITools/wolves.html, and we briefly outline those procedures here again. Data generated by Illumina sequencing were processed using OBITools (https://pythonhosted.org/OBITools/) command ‘ngsfilter’ to assign each sequence record to the corresponding sample based on tag and primer. Then ‘obiuniq’ was used to dereplicate reads into unique sequences. Reads less than 190 bp and with counts less than 10 were discarded. Denoising was performed using ‘obiclean’ to retain only sequences with no variants containing a count greater than 5% of their own. Sequences were assigned to taxa using ‘ecotag’ and a result table was generated using ‘obiannotate’. Our reference database was built in silico using our universal fish primer assay on 03/08/2021. Only fish species with identities ≥ 90% and whose sequence variants could be assigned to at least family (and lower) were included. All variants were assigned a single name (eg. to family, genus or species) and directly compared to the known species in the mesocosm (Table 2). For example, an assignment to genus could be compared to the species of that genus which are known to inhabit the mesocosm.

**Table 2.**
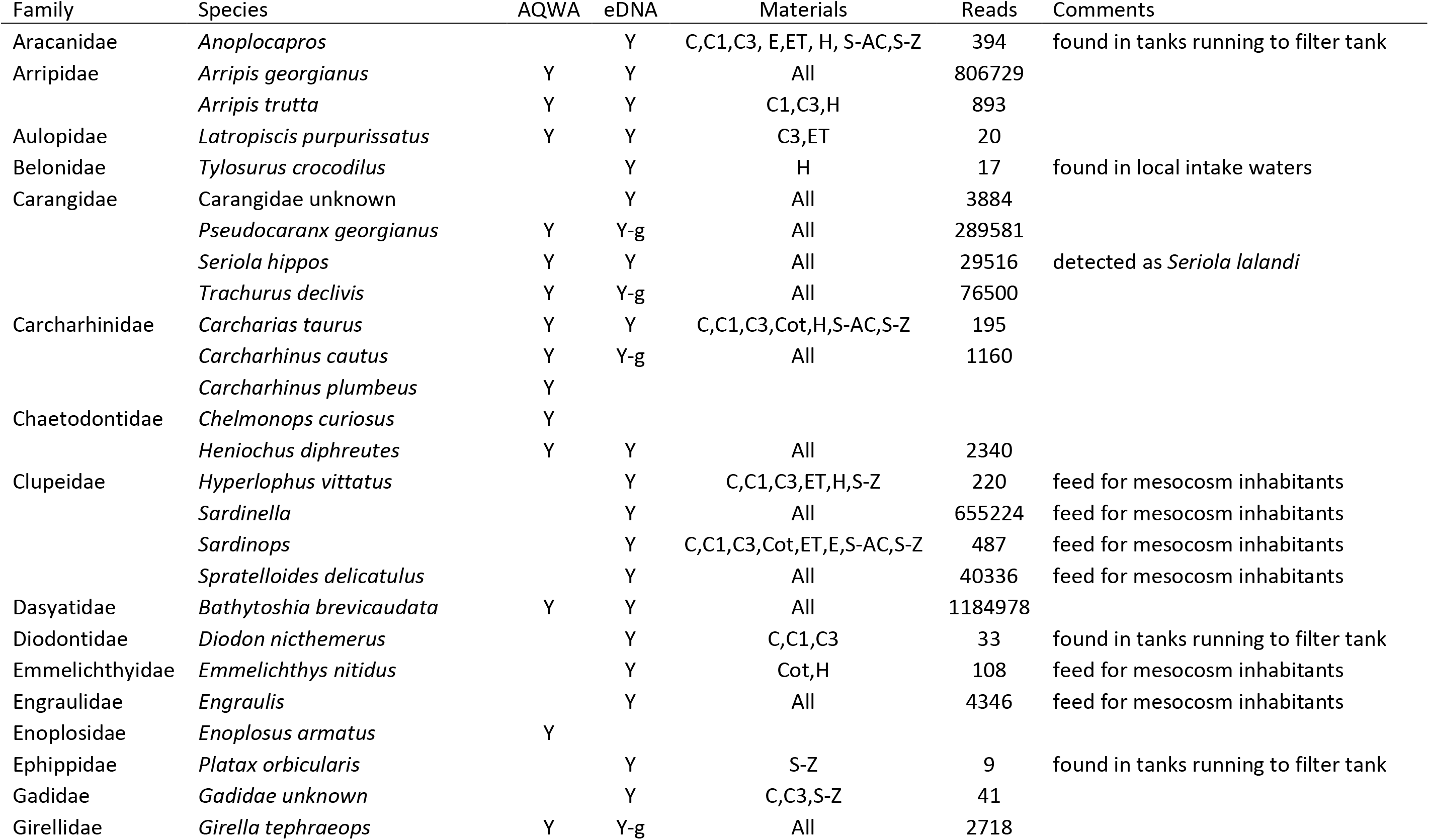
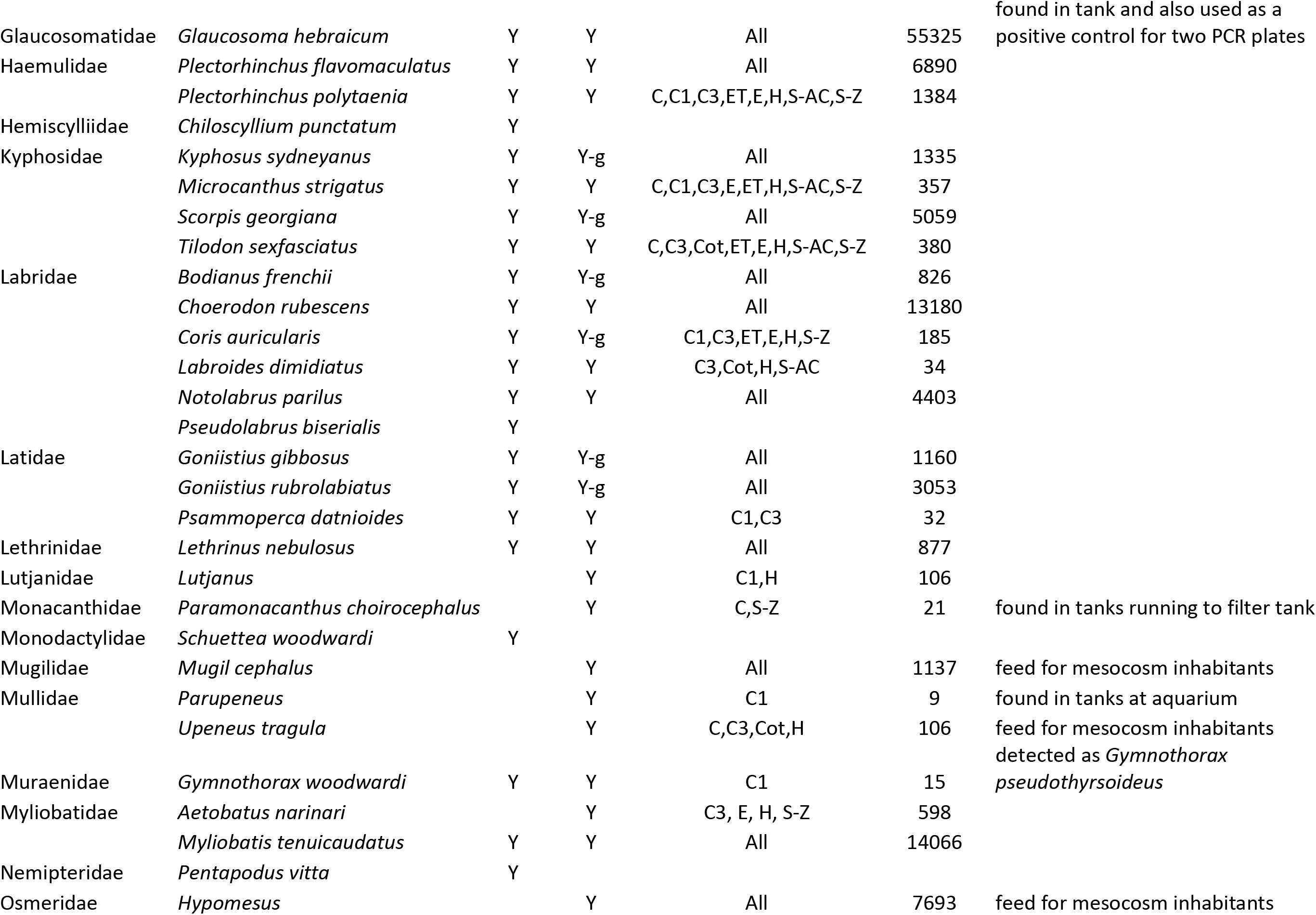
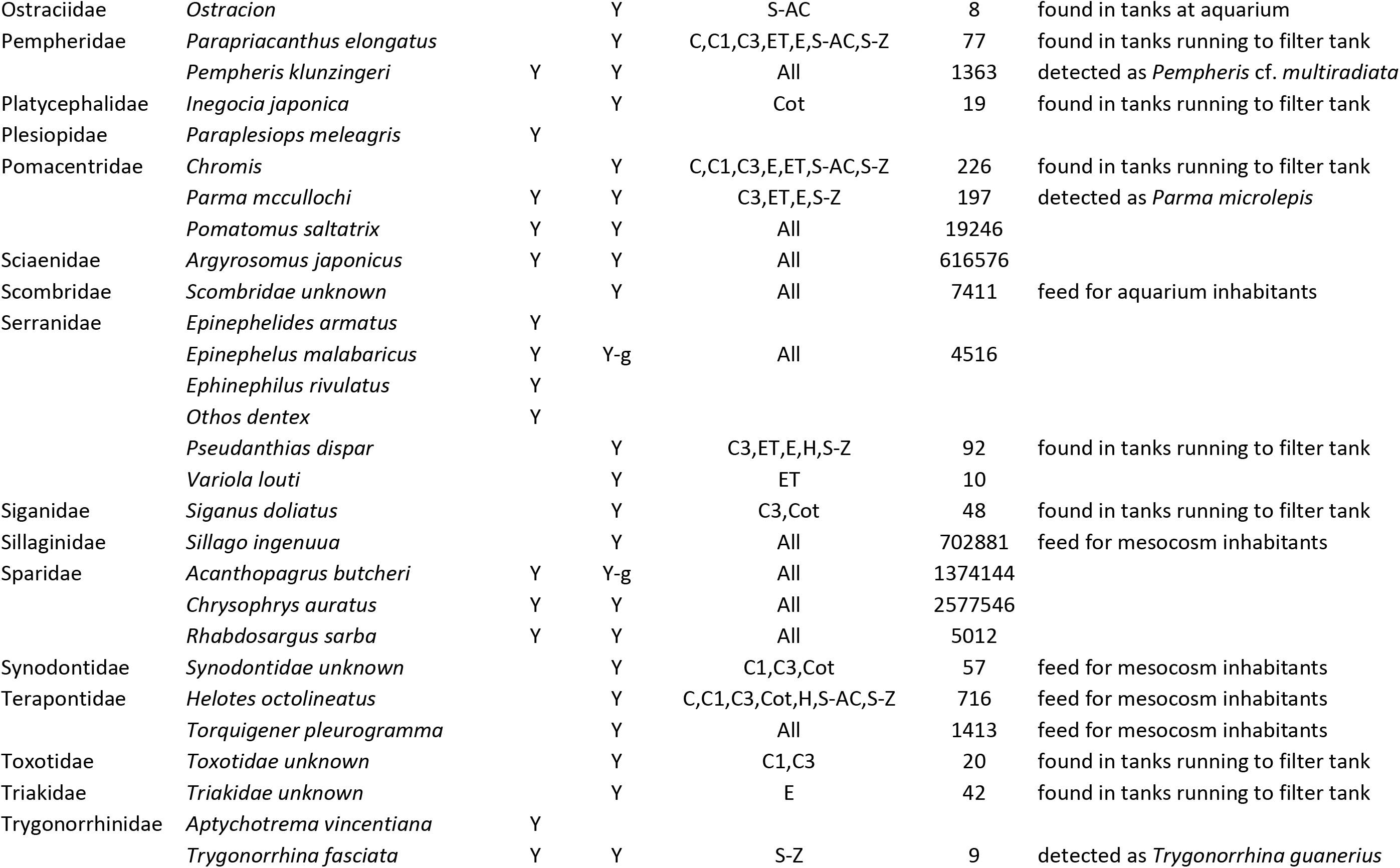
Fish species inhabiting the main tank compared to those detected by passive eDNA collection, the materials upon which each species were detected (C=cellulose, C1=chitosan-1%, C3=chitosan-3%, E=espun, ET=espun w/ chitosan, Cot=cotton, H=hemp, S-AC=sponge-active carbon, S-Z=sponge-zeolite), the number of reads and associated comments per species (Y-g=yes to genus).

### Statistics

A Box-Cox transformation normalized the data (Shapiro-Wilks Test), which allowed for the use of parametric statistics. We used an analysis of variance on the linear model fit of mean Cq value by material, followed by a Tukey Honest Significant Difference to compare materials. We also used an analysis of variance on the linear model fit between mean Cq value and submersion duration for each material. A linear model fit of mean Cq values by material and submersion duration, and their interactions, produced the same results. These statistics were likewise used to determine differences in the number of species detected between materials and submersion intervals. We fit a smoothing spline to the interval data for a visual estimation of how mean Cq values and species detections varied with time. All statistics and graphics were produced using R (version 2.14.0; R Development Core Team 2011) and graphics were edited in Inkscape (https://inkscape.org/).

## Results

### Multiple Materials Enable Passive eDNA Collection

All nine membrane materials collected detectable fish eDNA (Fig. 2). Although significant differences in mean Cq values existed between materials (F = 21.69, df = 11; p < 0.001), with cotton and hemp fibres exhibiting higher mean Cq values, all other materials were similar to each other, including to those obtained from conventionally filtering five 1L samples. Both cotton and hemp were deployed within nylon bags, and the mean Cq values for the nylon bags (22.3, 23.1, 23.7; min, mean, max) were not significantly different than that of the filtered 1L samples (21.1, 23.5, 28.5), nor most other materials. Additionally, the mean Cq values of the nylon bags were lower than that of the zeolite sponge (p = 0.01). The backing plate attached to the electrospun nanofibre covered cellulose membranes did not inhibit eDNA collection, as evidenced by their comparable mean Cq values.

**Figure 2.**
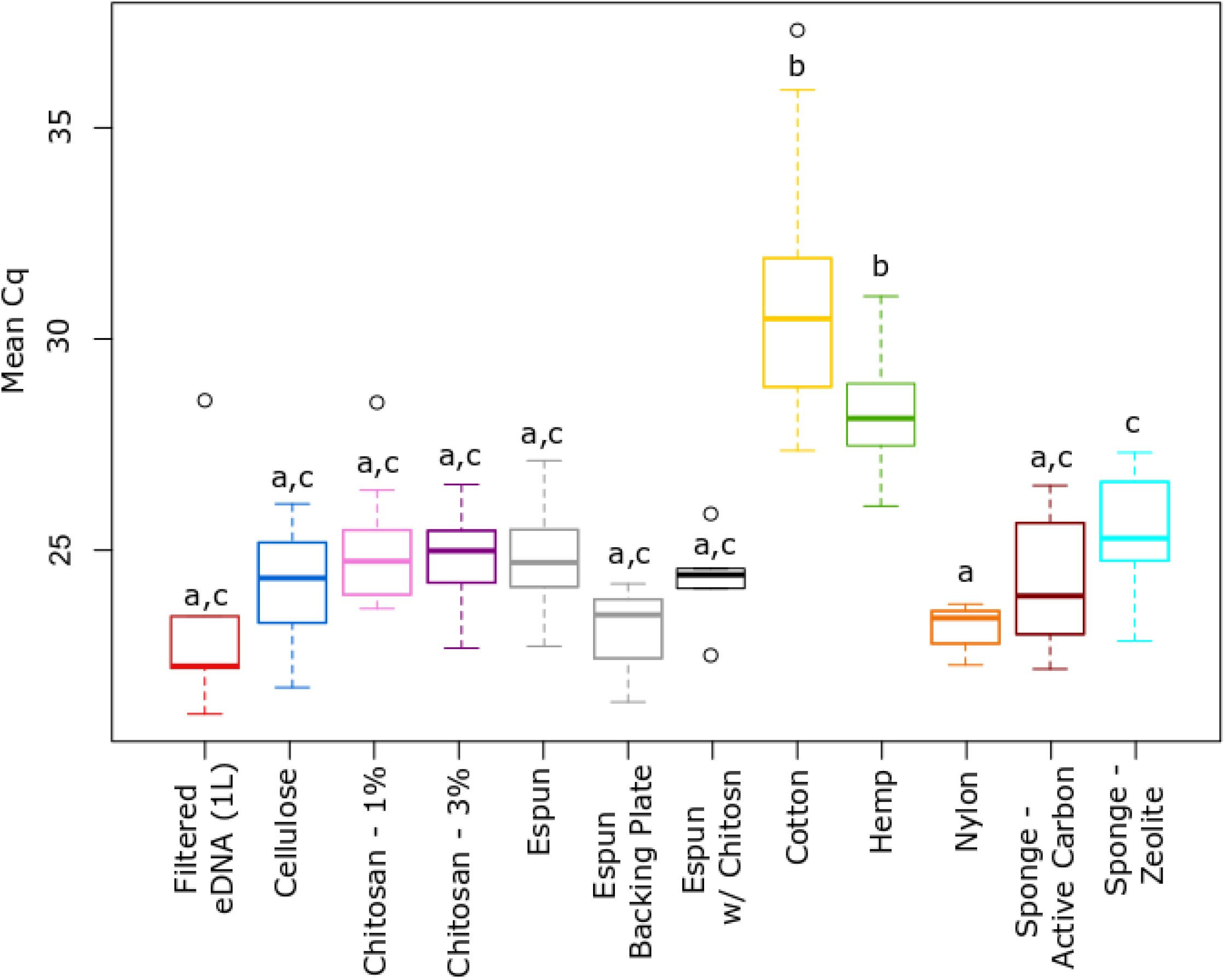
Mean Cq values from quantitative PCR by membrane material compared to conventional filtration of 1L eDNA samples. Note that both cotton and hemp were place inside nylon bags. Lower Cq values indicate higher DNA yield and different letters indicate statistical significance (α=0.05).

### Increased Submersion Time Did Not Increase eDNA Collection

No significant differences in mean Cq values were detected over time for any of the nine trialled materials (Fig. 3; F = 1.28 for submersion time * material, df = 9, p = 0.25). Smoothing splines were fitted to the time interval data for each material, and a trend downward would be indicative that the material collected more eDNA over time. Only hemp and both sponge materials showed any decline in trendline over time.

**Figure 3.**
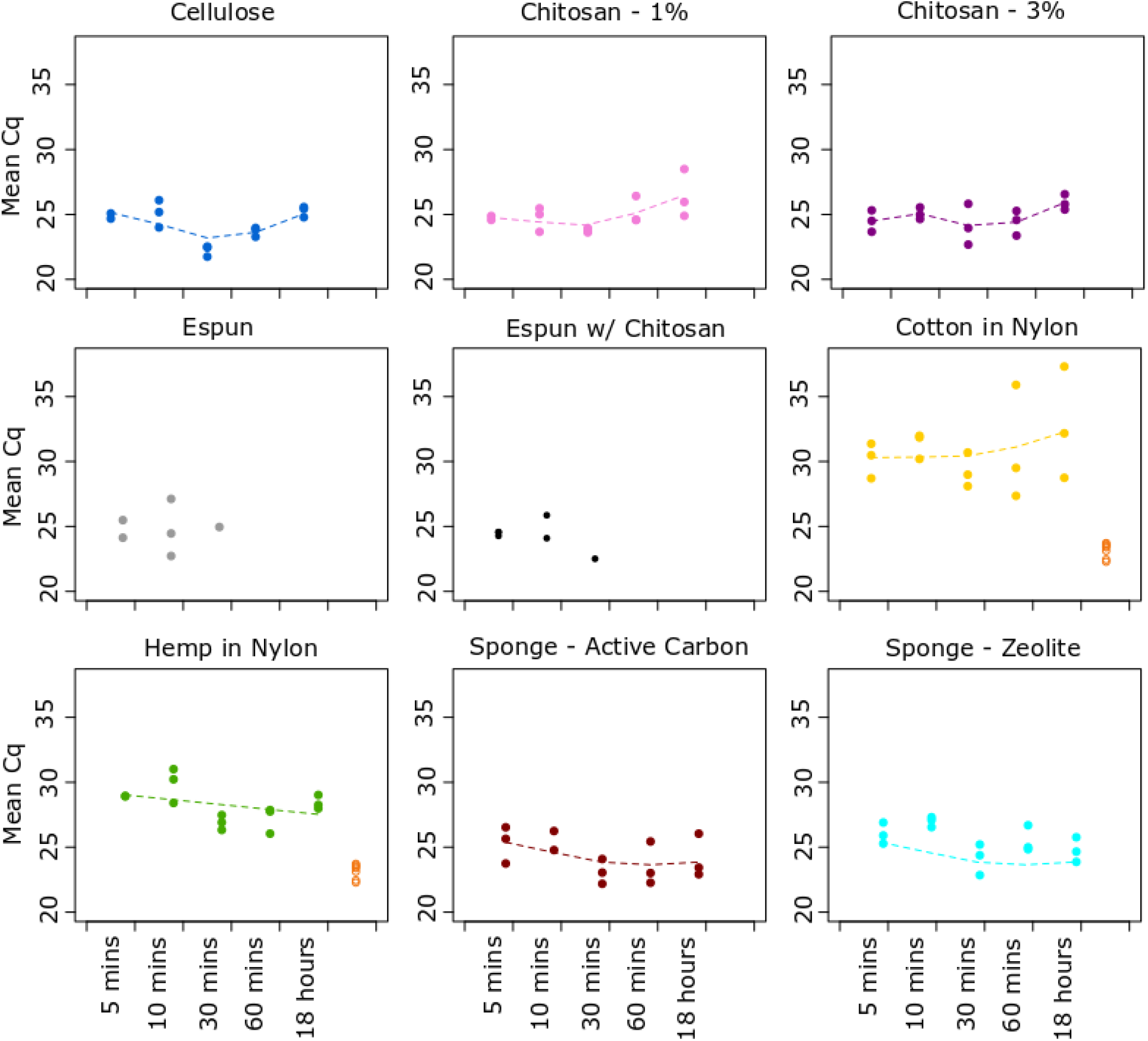
Mean Cq values from quantitative PCR by submersion time for each membrane material. Open circles represent data for nylon bags (used with both cotton and hemp) which were sampled only at the end of the experiment. A smoothing spline (dashed line) is used to visualize possible time trends and could not be fitted for electrospun nanofibers due to the small sample size.

### The Majority of Fish Species were Detected with All Materials

For all materials, we assigned 8,822,884 sequence reads to 71 fish taxa (Table 2). Of the 50 species known to inhabit the mesocosm, 37 (74%) were detected through passive eDNA. The additional 34 species detected include known feed taxa, fish found in local intake waters, and fish occupying tanks within the same facility and within the same water system. The number of species detected differed between materials (Fig.4; F=21.69, df=11, p < 0.01), with cotton, hemp and nylon detecting the fewest number of fish species on average. However, all materials detected a comparable number of species to the conventionally filtered eDNA samples. The median number of fish species detected by material was 27 (filtered eDNA; range = 19-32), 29 (cellulose; 19-33), 31 (chitosan - 1%; 17, 36), 31 (chitosan - 3%; 23, 37), 29 (espun; 22, 31), 26.5 (espun with chitosan; 21, 36), 19.5 (cotton; 11, 28), 24 (hemp; 14, 30), 24 (nylon bags; 18, 29), 27 (sponge – active carbon; 13, 37) and 29 (sponge – nitrate; 21, 34). The number of species detected did not differ with submersion time for any of the nine trialled materials (Fig. 5; F = 0.68 for submersion time * material, df = 9, p = 0.72).

**Figure 4.**
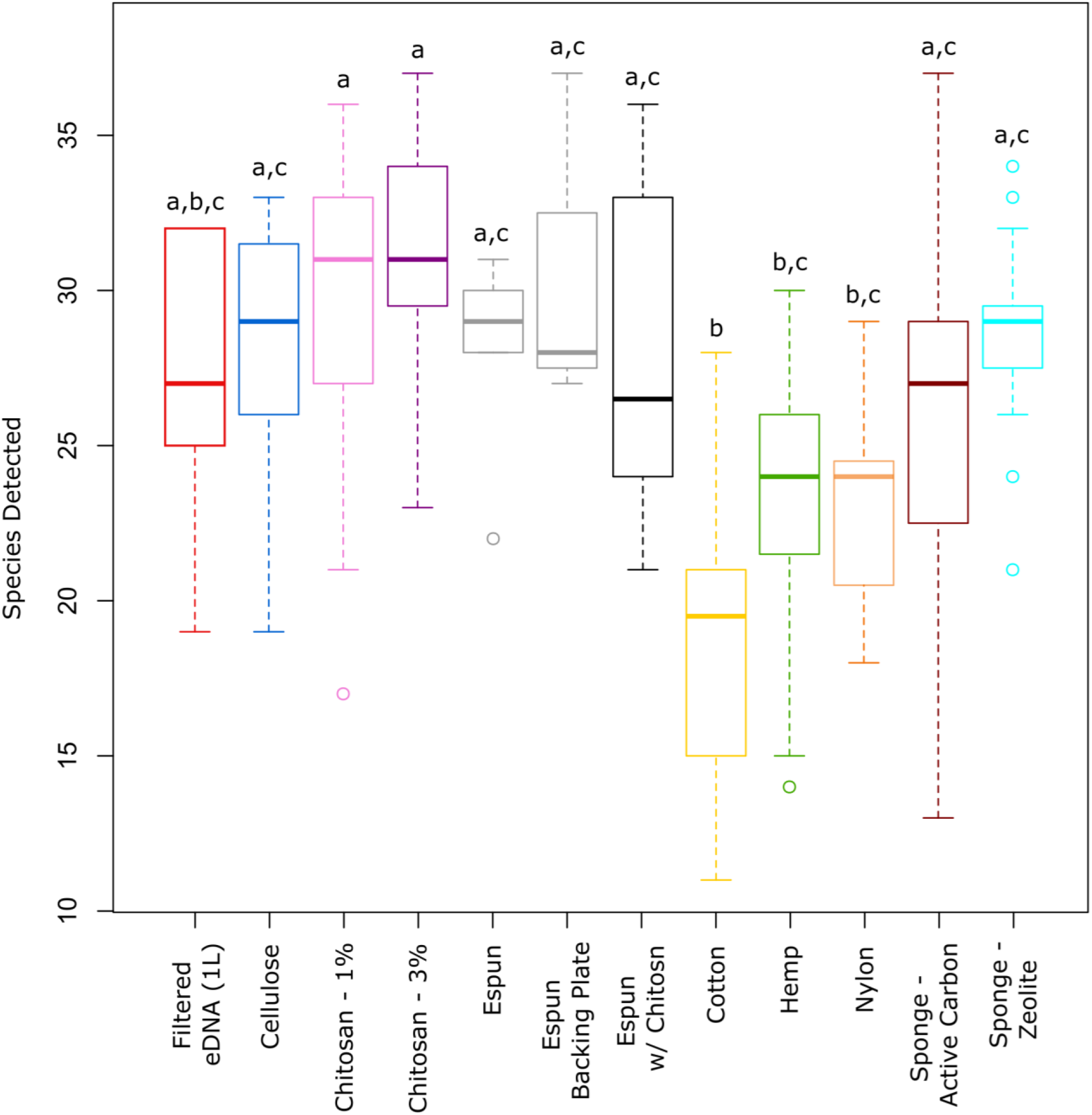
The number of fish species detected by membrane material compared to conventional filtration of 1L eDNA samples. Note that both cotton and hemp were place inside nylon bags. Different letters indicate statistical significance (α=0.05).

**Figure 5.**
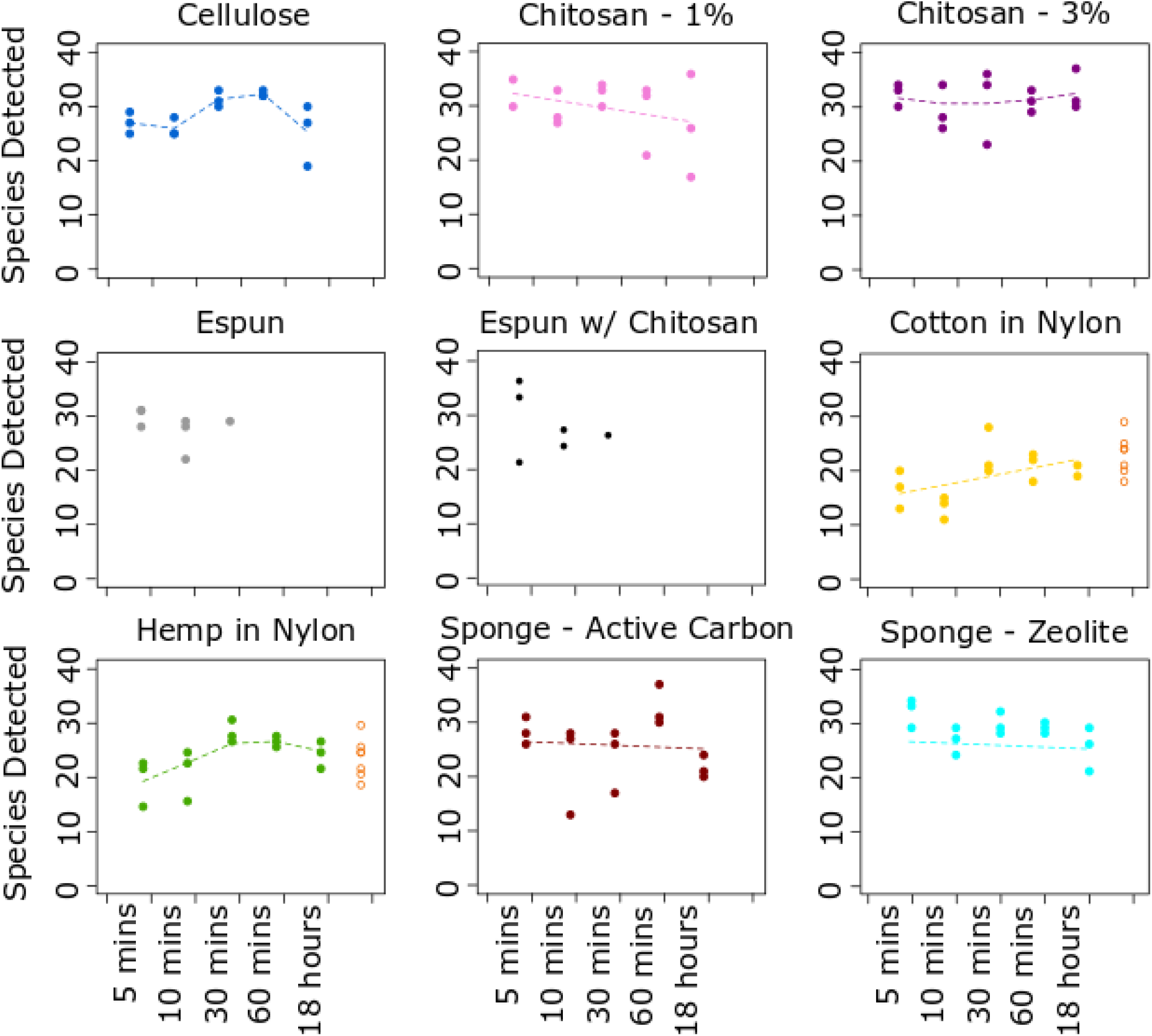
The number of fish species detected by submersion time and membrane material. Open circles represent data for nylon bags (used with both cotton and hemp) which were sampled only at the end of the experiment. A smoothing spline (dashed line) is used to visualize possible time trends and could not be fitted for electrospun nanofibers due to the small sample size.

### Scanning Electron Microscopy of eDNA Collection Materials

The SEM images displayed how biological matter was adhered to the surface of all material (Fig.6) and did not appear entrapped or bound in any consistent manner. These images also revealed the diversity in size and structure of biological matter found on the materials. For example, a ‘slick’ of biological material can be seen on some materials (see Fig.6 cellulose and chitosan – 3%) while others contain small, rounded particles of biological material (see Fig. 6 espun, cotton and hemp) or larger particles with irregular shapes that have a crystalline appearance or extremely smooth surface, consistent with inorganic materials and debris like salt crystals and sediment (see Fig.6 sponge – active carbon).

**Figure 6.**
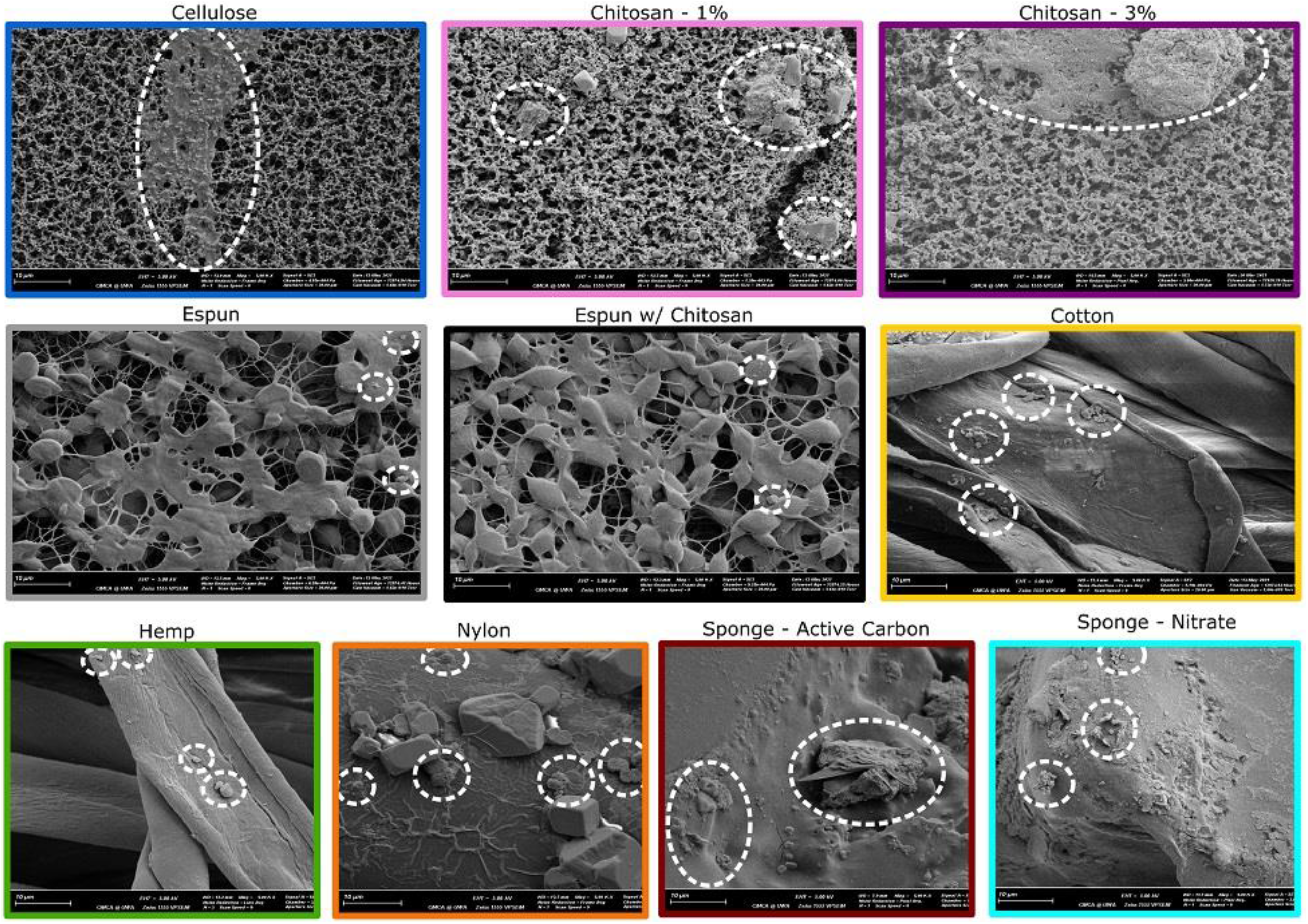
Scanning electron microscopy images of materials at 5000 × magnification after submersion in tank water where dashed circles identify biological matter. All scale bars are 10 μm.

## Discussion

We provide the first comprehensive evaluation of the capacity of a variety of porous materials to passively collect environmental DNA from a marine environment. Further, we also test the importance of submersion time. We reveal that numerous inexpensive materials are highly effective for the passive collection of eDNA. Remarkably, we also show that passive eDNA collection can be as effective as conventional water filtering and achieved quickly; in as little as five minutes.

## Materials

We identify multiple materials that can be used effectively for passive eDNA collection. Material choice can influence capture efficiency, and ideally, the selected material will maximize eDNA capture without interfering or complicating the extraction process. We investigated the use of materials which varied in structural complexity and robustness, and found no significant difference in capture efficiency, apart from the reduced capacity of hemp and cotton fibres that were contained in nylon bags. Kirtane et al. (2020) likewise found that the pore size used to encase the adsorbent had a significant impact on DNA adsorption. They suggest that restricted flow over the adsorbent was associated with smaller pore sizes, and that increasing encasing pore size increases capture. These results highlight the importance of membrane encasing and suggest the maximum surface area of a material should come in direct contact with the water.

There was also some indication that the addition of chitosan to a material could increase capture efficiency, since both chitosan treatments detected the highest median and maximum fish species richness. Although this was not statistically significant, our experiment was conducted in a low diversity, temperate mesocosm. Since water characteristics, such as pH and temperature, can influence DNA adsorption to different materials (Lorenz and Wackernagel 1987), it’s possible the addition of chitosan could result in increased capture efficiencies in some environments. For example, Bessey et al. (2021) found that nylon membranes performed as well as conventional filtering for fish species detection in low-diversity temperate waters but not in high-diversity tropical waters, presenting a situation where the addition of chitosan could potentially increase the effectiveness of nylon materials. Further investigation into material optimization will be particularly important for high diversity systems.

Practical considerations for material choice will play an important role since our results reveal many materials are effective for passive eDNA collection. Cost of material, availability, robustness, ease of deployment and downstream processing, may all influence material choice. For example, nanomaterials are more time-consuming and costly to produce than readily available cellulose ester membranes and aquarium grade sponges, which are commercially available (Liu 2012). Cellulose ester membranes require less handling time during downstream processing than granular materials, which require weighting, or sponge and nylon materials that require cutting prior to DNA extraction (Bessey et al. 2021). However, in a turbulent, high flow water environment, a more robust material, such as sponge, may be desirable over the more fragile cellulose membranes. A challenge for employing passive eDNA collection will be finding a standard that can be consistently used so that time series and spatial comparisons are meaningful within and between studies. This challenge similarly exists for conventional eDNA studies (Trujillo-Gonzalez et al. 2021).

### Time

We determined that long submersion times are not necessary for passive eDNA collection and effective sampling can be achieved in as little as five minutes. Conventional eDNA filtration methods are time consuming, especially when considering the amount of water needed to effectively filter an area for accurate biodiversity estimates (Bessey et al. 2020, Koziol et al. 2019). A quick eDNA collection method would have considerable benefit for end users and increase sampling capacity. Increased sampling capacity enables a broader range of ecological question to be addressed through comparative frequency analysis (Strickland et al. 2019), which are more powerful with larger sample sizes. Therefore, exploring the minimum amount of time required for passive eDNA collection membranes to saturate would be a worthwhile endeavour to maximize efficiency. In laboratory experiments, Kirtane et al. (2020) found no difference in adsorbed extracellular DNA concentrations over a time gradient between one min to two hours for granular active carbon in tank water, but that differed in creek waters. These previous studies indicate that site specific water chemistry affects the effectiveness of passive eDNA collection. Therefore, a better understanding of the mechanism of eDNA adherence to materials could help optimize passive eDNA collection methods.

### Mechanism of eDNA Adherence to Materials

The mechanism by which eDNA adheres to materials in natural aquatic systems remains unclear. We used scanning electron microscopy to gain insight into the mechanism of attachment but found no consistent patterns. Despite trialling materials with a range of surface complexities, we found no supporting evidence that eDNA was entrapped within the interstitial spaces of the materials. Rather, biological matter appeared to adhere randomly to any available surface and showed great diversity in size and shape. For example, morphologically distinct single cell eukaryotes and bacteria could be seen on the surfaces of the membrane materials, many embedded in larger bodies of seemly biological material, most likely biofilm. An important component of biofilm development is extracellular polymeric substances (Hancock 2001, Vilain et al. 2009) which are mainly comprised of polysaccharides, proteins, metabolites and extracellular DNA (Das et al. 2013). These extracellular polymeric substances occur in a range of molecular sizes, conformations and physical/chemical properties and although little is known about the physical ultrastructure of how they interact (Decho and Gutierrez 2017), they are known to adhere to both natural and engineered surfaces (Das et al. 2013). The diversity of biological compounds and structures that eDNA might be associated with in aquatic systems is huge. Dissolved organic matter (DOM) may contain more than 20,000 compounds in a single seawater sample (Mentges et al. 2017). Particulate organic matter (POM) as seen in Figure 6 contains equal or greater diversity as well as structural complexity because much of it is derived from dead organisms (Kharbush et al. 2020). A deeper understanding of the adhesive properties of different fractions of the POM pool and biofilms associated with passive eDNA collection materials may provide deeper insights into eDNA binding to collection materials.

### Future Research

We contribute to building evidence that passive eDNA collection is effective and offers important advantages over conventional water filtration methods, warranting further investigation. Studies conducted in environments where eDNA degrades quickly or is released in pulses may identify further advantages of passive eDNA collection. Although our study identified both rare and abundant species in a relatively low diversity mesocosm, future studies should evaluate the effectiveness of materials and submersion time in warmer, high diversity systems so that materials are effective for the maximum number of environmental conditions. Even in situations where passive eDNA collection may not perform as optimally as conventional filtering methods, the time and cost efficiencies may still warrant it’s use, making cost-benefit analysis of which method to use a worthwhile consideration. Futures studies focussed on a mechanistic understanding of not only how eDNA adheres to materials, but investigating the physical/chemical properties of eDNA, could lead to the greatest advances in passive eDNA collection methods and optimization of materials.

## Data Availability

Raw sequences, bioinformatic script, reference database, and the final datasets are available on the CSIRO Data Access Portal at xxx (raw sequence and final dataset) and https://data.csiro.au/collections/collection/CIcsiro:46025v1 (bioinformatic script and reference database).

## Acknowledgments

This project was funded by the CSIRO Environomics Future Science Platform. We thank Matt O’Malley and AQWA for allowing our experiments to be conducted at the facility. The authors acknowledge the facilities, and the scientific and technical assistance of the Australian Microscopy, Characterisation & Analysis, the University of Western Australia, a facility funded by the University, State and Commonwealth Governments. The authors also thank Sylvia Osterrieder and Zoe Slatter for their invaluable help during the mesocosm experiment and in the laboratory.

## Author Contributions

CB, YG, YT, HM, SJ, OB– contributed intellectual direction

CB– designed study

YG, YT – designed membrane materials

HM – conducted SEM imaging and interpretation

CB – organized/participated in mesocosm sampling

CB– performed molecular research

CB – processed and analysed all data

CB – conducted statistical analysis, produced graphics and tables

CB, YG, YT, HM, SJ, OB– assisted with manuscript writing

All authors contributed to manuscript revisions

